# Methyl Carnosate, a Carnosic Acid Derivative, Attenuates Osteoclastogenesis via Modulation of RANKL-Induced NF-κB Activity

**DOI:** 10.64898/2026.01.15.699687

**Authors:** Lifang Zhang, Chengxu Xie, Xinyi Bao, Mojtaba Tabandeh, Vishwa Deepak

## Abstract

Excessive osteoclast activity underlies bone-destructive diseases including osteoporosis and rheumatoid arthritis. RANKL-induced NF-κB signaling represents a critical pathway driving osteoclastogenesis. Here, we report that methyl carnosate, a naturally occurring diterpene from rosemary, inhibits RANKL-induced osteoclastogenesis with an IC_50_ of 1.2 μM and displays greater potency than its parent compound, carnosic acid. Promoter–reporter analysis further indicates attenuation of RANKL-induced NF-κB activity. These findings identify methyl carnosate as a potential lead compound for the development of bone-protective therapeutics.

## INTRODUCTION

Dysregulated osteoclast activity drives bone loss in osteoporosis, inflammatory arthritis, and bone metastases (Takegahara et al. 2024). RANKL-induced NF-κB signaling is essential for osteoclast differentiation, making it an attractive therapeutic target (Zhou et al. 2022). Current anti-resorptive agents have limitations including adverse effects, highlighting the need for novel therapeutics (Naciu et al. 2025).

Carnosic acid, a phenolic diterpene from rosemary, inhibits RANKL-induced osteoclastogenesis at concentrations of 5–15 μM. However, the relatively high effective doses and limited stability may constrain its therapeutic potential (Thummuri et al. 2017; Zheng et al. 2020). Methyl carnosate, a naturally occurring methyl ester derivative of carnosic acid, exhibits enhanced antioxidant activity and stability in lipid systems (Climati et al. 2013; Huang et al. 1996) . However, its effects on osteoclastogenesis remain unexplored. Given that esterification typically enhances cellular permeability, we hypothesized that methyl carnosate might exhibit improved anti-osteoclast potency. Here we investigated whether methyl carnosate inhibits RANKL-induced osteoclast differentiation and assessed its mechanism of action.

## MATERIALS AND METHODS

### Reagents and Cell Culture

Methyl carnosate (CAS 82684-06-8, purity ≥98%, MedChemExpress, NJ, USA) was dissolved in DMSO (10 mM stock, final concentration ≤0.1%). RAW 264.7 murine macrophages were cultured in DMEM with 10% FBS at 37°C, 5% CO_2_. For osteoclast differentiation, cells were treated with 50 ng/mL recombinant murine RANKL (R&D Systems, MN, USA) ± methyl carnosate for 5 days.

### Cell Viability and TRAP Activity Assays

Cell viability was assessed using CCK-8 assay (0.1-10 μM methyl carnosate, 48-72 h). TRAP activity was measured using a colorimetric kit (#EEA055, ThermoFisher Scientific, MA, USA) following osteoclast differentiation. IC_50_ values were calculated using GraphPad Prism 9.

### NF-κB Reporter Assay

RAW 264.7 cells stably expressing NF-κB-luciferase reporter (#D2206, Beyotime, China) were pre-treated with vehicle or methyl carnosate (1 h), then stimulated with RANKL (6 h). Luciferase activity was measured using Steady-Glo® system (Promega, WI, USA).

### Statistical Analysis

Data represent mean ± SD from ≥3 independent experiments. Statistical significance was determined by one-way ANOVA with Tukey’s post-hoc test (*p* < 0.05 considered significant).

## RESULTS AND DISCUSSION

Methyl carnosate (Figure 1A) maintained >95% cell viability at ≤5 μM in RAW 264.7 macrophages at 48 and 72 h, with viability >80% even at 10 μM (Figure 1B). The compound dose-dependently inhibited RANKL-induced TRAP activity with an IC_50_ of 1.2 μM (Figure 1C), demonstrating approximately 4-12-fold greater potency than carnosic acid (5-15 μM) (Thummuri et al. 2017; Zheng et al. 2020). This substantial improvement likely reflects enhanced cellular permeability from increased lipophilicity conferred by the methyl ester. The IC_50_ lies well below cytotoxic concentrations, indicating a favorable therapeutic window and specific pathway inhibition rather than general cytotoxicity.

**Figure 1.**
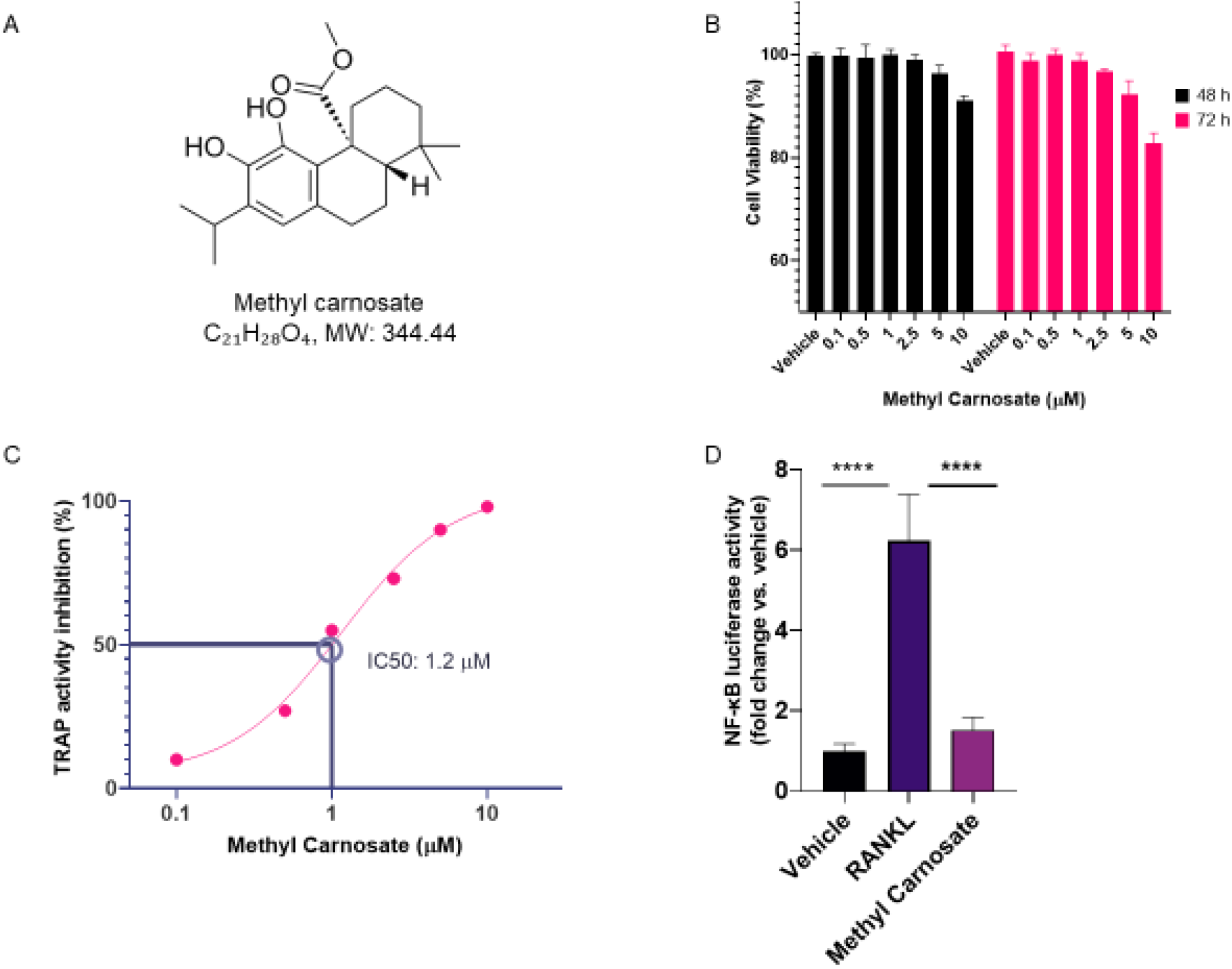
Methyl carnosate inhibits RANKL-induced osteoclastogenesis through NF-κB pathway suppression. (A) Chemical structure of methyl carnosate. (B) Cell viability of RAW 264.7 cells treated with indicated concentrations of methyl carnosate for 48 hours (black bars) or 72 hours (pink bars), measured by CCK-8 assay. (C) Dose-dependent inhibition of TRAP activity in RANKL-stimulated RAW 264.7 cells. IC50 = 1.2 μM. (D) Effect of methyl carnosate on RANKL-induced NF-κB transcriptional activity measured by luciferase reporter assay. Data represent mean ± SD from three independent experiments. Statistical significance was determined by one-way ANOVA (\*\*\*\**p* < 0.0001 vs. RANKL alone).

To elucidate the mechanism, we examined NF-κB signaling, a critical early mediator of RANKL signaling. Using an NF-κB-luciferase reporter assay, methyl carnosate significantly suppressed RANKL-induced NF-κB transcriptional activity (Figure 1D). RANKL stimulation induced robust NF-κB activation (∼6-fold over control), whereas methyl carnosate pre-treatment reduced this to near-basal levels, demonstrating interference with NF-κB signaling essential for initiating the osteoclastogenic program (Asagiri and Takayanagi 2007).

The markedly enhanced potency of methyl carnosate compared to carnosic acid provides important structure-activity insights. Esterification yields increased lipophilicity enhancing membrane permeability, improved metabolic stability (Huang et al. 1996), and reduced susceptibility to oxidative degradation. These modifications result in >4-fold improvement in anti-osteoclast potency, positioning methyl carnosate as a superior lead compound.

Our findings establish that methyl carnosate inhibits osteoclastogenesis through NF-κB suppression. Future studies should examine effects on additional RANKL pathways, validate efficacy in primary bone marrow macrophages, and evaluate bone-protective effects in animal models.

## CONCLUSIONS

Methyl carnosate potently inhibits RANKL-induced osteoclast differentiation by suppressing NF-κB signaling, exhibiting significantly greater activity than its parent compound carnosic acid. These findings identify methyl carnosate as a promising lead compound for developing novel therapeutics targeting pathological bone resorption.

## ACKNOWLEDGMENTS

This work was supported by the WKU-ISRG grant (ISRG2023032) and the International Frontier Interdisciplinary Research Institute of Wenzhou-Kean University (IFIRI-WKU) grant (KY20250604000450) awarded to Vishwa Deepak, and by the WKU-ISRG grant (ISRG2023027) awarded to Mojtaba Tabandeh. We thank the core facilities of Wenzhou-Kean University for providing the necessary resources.

